# Analyzing interaction of rhodacyanine inhibitor ‘MKT-077’ with *Plasmodium falciparum* HSP70 nucleotide binding domains

**DOI:** 10.1101/2023.07.27.550796

**Authors:** Chanchal Nainani, Vipul Upadhyay, Bikramjit Singh, Komalpreet Kaur Sandhu, Satinder Kaur, Rachna Hora, Prakash Chandra Mishra

## Abstract

MKT-077 and its derivatives are rhodacyanine inhibitors that hold potential in treatment of cancer, neurodegenerative diseases and malaria. These allosteric drugs act by inhibiting the ATPase action of heat shock proteins of 70kDa (HSP70). MKT-077 accumulates in the mitochondria and displays differential activity against HSP70 homologs. The four *Plasmodium falciparum* HSP70s (PfHSP70) are present at various subcellular locations to perform distinct functions. In the present study, we have used bioinformatics tools to understand interactions between MKT-077 and PfHSP70s. Nature of the identified interactions is primarily hydrophobic with different PfHSP70 homologs showing variable propensity to bind MKT-077. Our molecular docking approach has helped us to predict the binding pocket and specific residues on PfHSP70s that are involved in interacting with MKT-077. Information derived from this article may form the foundation for design and development of MKT-077 based drugs against malaria.

## Introduction

*Plasmodium falciparum* (Pf) caused malaria remains a leading cause of death in tropical regions of the globe [1]. This pathogenic protozoan expresses a set of four heat shock proteins of 70 kDa (PfHSP70) that help the parasite in acclimatizing to stress conditions including temperature variations [2]. Different PfHSP70s localize variably to the parasite nucleus and cytoplasm (PfHSP70-1), endoplasmic reticulum (PfHSP70-2), mitochondria (PfHSP70-3) and cytoplasm/ J dots present in parasitized erythrocytes (PfHSP70-x) [3]. HSP70s are chaperones that have roles in folding, refolding, transport and targeting of cellular proteins through ATP hydrolysis dependent cycles of substrate binding and release [4]. HSP70s have a conserved domain architecture with their N-terminal region comprising the nucleotide binding domain (NBD) that performs ATPase activity. ATP hydrolysis to ADP is augmented by binding of HSP70 (via their NBD) with diverse co-chaperones ‘HSP40’. HSP70s are subject to allosteric regulation where ATPase action (ADP bound state) facilitates substrate binding of HSP70 via its C-terminal substrate binding domain (SBD). On the contrary, ATP binding to NBD significantly diminishes affinity of HSP70 for its client proteins. The ATP and ADP bound states of HSP70 are structurally distinct. While the substrate bound ADP form has the nucleotide binding cleft in its ‘open’ state, this surface cleft is ‘closed’ in the apo ATP bound form of HSP70.

The cellular levels of various HSP70s are upregulated in many cancers to manage stress, prevent apoptosis and promote cell survival [5]. These act by blocking cell death pathways, directly inactivating the tumor suppressor ‘p53’ and possibly negating structural changes in mutant proteins expressed by cancerous cells. Therefore, inhibition of HSP70s is being exploited as a possible anti-cancer mechanism. HSP70 inhibitors have also shown promise in the treatment of neurodegenerative diseases like Alzheimer’s *etc*. which are characterized by buildup of tau tangles. MKT-077, a synthetic rhodacyanine derived HSP-70 inhibitor that accumulates in the mitochondria is reported to preferentially inhibit growth of five different cancerous cell lines (IC_50_: 0.35 – 1.2 uM) when compared with normal epithelial cells (100-fold higher IC_50_ values). MKT-077 also helps in clearance of tau angles, but it is unable to cross the blood brain barrier and hence considered unsuitable for treatment of central nervous system disorders [6]. This molecule showed significant promise to be developed as an anti-cancer drug owing to its favorable characteristics like solubility, stability, toxicity and pharmacokinetics in pre-clinical trials. Although MKT-077 was unable to pass a Phase I trial owing to renal toxicity [7], its pharmacological and toxicological parameters were found acceptable in another Phase I trial where lower doses were administered [8]. Despite a few shortcomings of MKT-077 like short half-life, there is continued interest in this molecule and its derivatives for development of cancer therapeutics. Rhodacyanine dyes have also been reported to be effective in selectively killing cultured Pf parasites with EC_50_ values ranging from 3-400 nM [9]. In this study, MKT-077 was found to be strongly active (EC_50_ = 70 nM) with good selective toxicity of 210.

The chemical formula for MKT-077 is C_21_H_22_N_3_OS_2_Cl (l-ethyl-2-{[3-ethyl-5-(3-methylbenzothiazolin-2-yliden)]-4-oxothiazolidin-2-ylidenemethyl) pyridium chloride) with a molecular weight of 432.01. It is a highly water-soluble molecule (>200 mg/ml). MKT-077 binds to the NBD of HSP70 and inhibits its ATPase activity owing to the induced conformational changes [10]. It is also reported to accumulate in mitochondria and target mortalin, the constitutively expressed mitochondrial HSP homolog in humans (also called HSPA9). Mortalin is likely to be involved in mitochondrial protein import through TIM (Translocase of Inner Membrane) [11-14]. MKT-077 destabilizes the ATP bound state of mortalin and weakens its p53 suppression activity resulting in complement-mediated cancer cell death [14-16]. MKT-077 is also known to bind to an allosteric site in NBD of the cytosolic human homolog (Hsc70/ HSPA8) and lock the molecule in a pseudo-ADP bound conformation. Since MKT-077 binds to the open and closed allosteric states of HSP70 differentially, it is considered to be an allosteric drug. The binding pocket of MKT-077 on HSPA8 has been identified through nuclear magnetic resonance (NMR) [5]. In the current study, we have used an *in-silico* approach to identify the binding pocket of MKT-077 on PfHSP70s. Binding pocket analysis of different PfHSP70 homologs has helped us to identify their specific amino acid residues that may be involved in MKT-077 interaction, and may form the foundation for development of new antimalarials.

## MATERIALS AND METHODS

### Obtaining sequences and structures

All four PfHSP70 homologs have been analyzed for their binding with MKT-077 in the present study (Table 1). Structures of NBDs of PfHSP70-2 and PfHSP70-x are solved experimentally, and were hence downloaded from PDB database (5UMBand 6S02 respectively) [17]. Sequences of PfHSP70-1 and PfHSP70-3 were downloaded from plasmodb.org and used for homology modeling by SwissModel [18] [19]. The obtained models were validated by Ramachandran plot analysis and ERRAT before using for further investigation and docking experiments. Structure of the inhibitor MKT-077 was downloaded from PubChem database in SDF format.

**Table 1:** The PfHSP70 homologs and their PlasmoDB IDs are tabulated.

| **HSP70 homologs** | **Plasmodb ID** |
| --- | --- |
| PfHsp70-1 | PF3D7_0818900 |
| PfHsp70-x | PF3D7_0831700 |
| PfHsp70-2 | PF3D7_0917900 |
| PfHsp70-3 | PF3D7_1134000 |

### Identification of probable binding region for MKT-077

We have explored two binding sites on PfHSP70s for the binding of MKT-077. One is the ADP binding site based on the crystallographic structure of PfHSP70-2 NBD complexed with ADP (5UMB) and a neighboring binding site obtained from a solved structure of PfHSP70-x complexed with HEW (2-amino 4 bromopyridine) (7OOG). These structures were downloaded from PDB and analyzed using LigPlot to identify the small molecule binding regions **[20]**. Binding regions for other homologs corresponding to both ADP and MKT-077 were predicted by using sequence alignment [21].

### Molecular docking and analysis

The protein structures and inhibitor were prepared for docking by using UCSF chimera where modifications like removal of non-standard residues, addition of H-bonds *etc*. were performed [22]. Based on the predicted ADP and HEW binding sites on PfHSP70s, gridlines were selected for docking of the inhibitor on all the structures. Following this, MKT-077 was docked on both the binding sites on all PfHSP70s through Autodock Vina plugin in Chimera [23]. Binding energies for each of the docked complexes were calculated in Chimera and the best docked complexes analyzed by using LigPlot.

## Results and discussion

### Preparation of protein and ligand structures for docking

MKT-077 is reported to bind with human counterparts HSPA8 and 9 of PfHSP70-1 and PfHSP70-3 respectively [14]. Here, we have tried to identify the binding sites for MKT-077 on various PfHSP70 homologs in an attempt to understand this interaction and further inhibitor discovery in this area. Since the structures of nucleotide binding domains of PfHSP70-x and PfHSP70-2 are reported, these were obtained from PDB database (6S02 and 5UMB respectively) (Fig 1a-b). Structures of the NBDs of the other two homologs i.e. PfHSP70-1 and PfHSP70-3 were obtained through homology modeling by using SwissModel [19] (Fig 1c-d). Structural integrity of the obtained models was ascertained through Ramachandran plot analysis and ERRAT (Fig S1). Structure of MKT-077 (SDF format) was obtained from PubChem (CID 6444403) (Fig 1e) [24].

**Figure 1:**
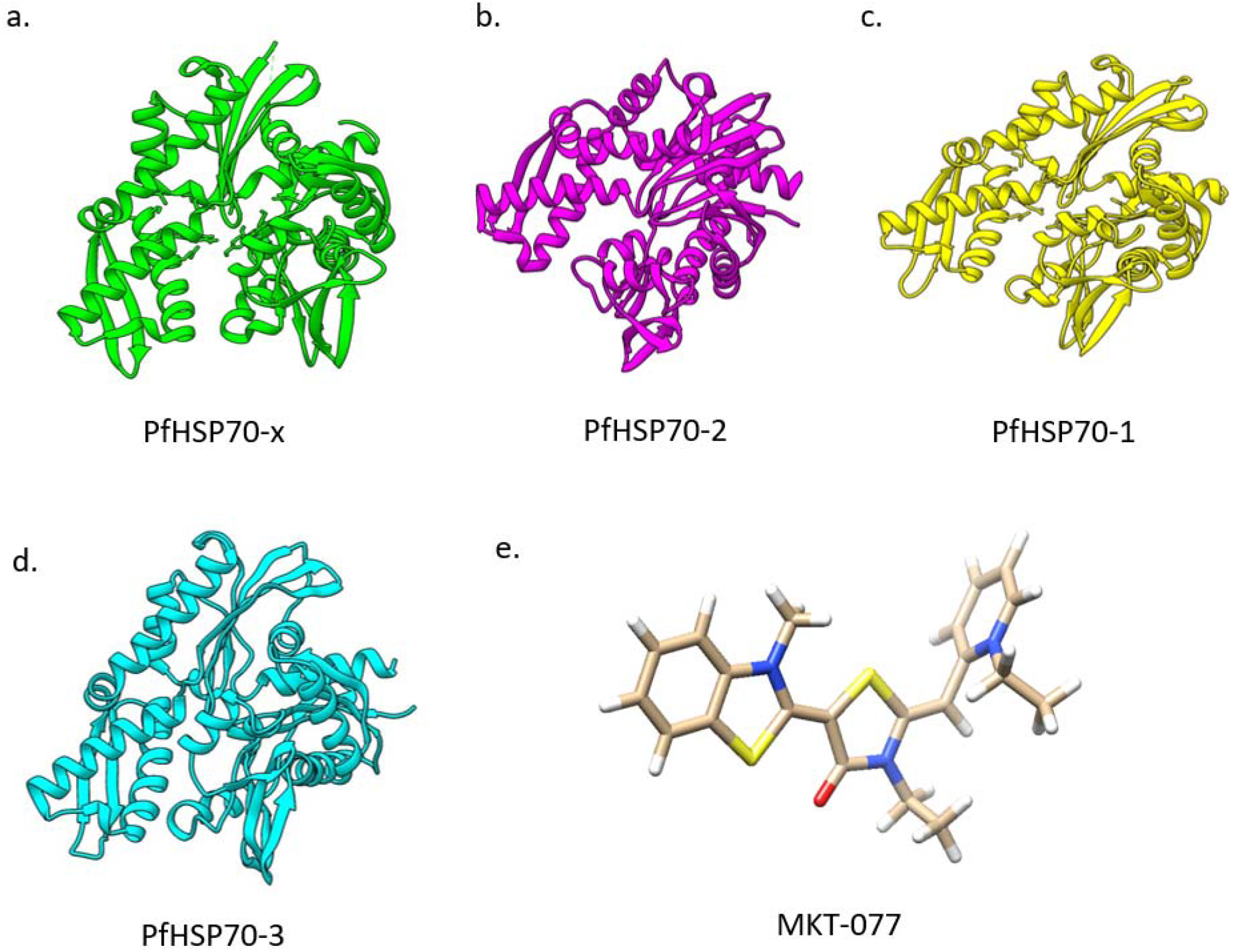
Structures of proteins and ligand. Cartoon representation of structures of nucleotide binding domains (NBD) of proteins obtained from PDB a. PfHSP70-x (6S02) b. PfHSP70-2 (7OOG). Protein structures of NBDs obtained through molecular modeling by SwissModel c. PfHSP70-1 and d. PfHSP70-3. e. Structure of MKT-077 obtained from PubChem (CID 6444403) in SDF format.

### Molecular Docking of MKT-077 on PfHSP70s

MKT-077 was docked on the prepared three dimensional structures of NBDs of all four PfHSP70s by using AutoDock Vina plugin in UCSF Chimera [22]. Docking experiments were performed on two distinct sites i.e. the ADP and HEW (an inhibitor) binding sites on NBD of PfHSP70s [25]. Binding site of ADP on PfHSP70-2 and HEW binding site of PfHSP70-x were identified on the basis of their solved three dimensional structures available on PDB (5UMB& 7OOG respectively) using LigPlot analysis [20] (Table 2). Binding sites for each of these ligands in the other homologs were predicted by using sequence alignment (Table 2). Grid boxes were selected on all the PfHSP70s based on the above binding region information to mark the interacting regions of proteins to be used for docking (Fig. 2). Coordinates for the grid boxes are given in figure S2 (Fig. S2). Results of molecular docking provided ten different poses of the docked complexes for each binding site.

**Table 2:** ADP and HEW binding residues on PfHsp70 homologs. ADP and HEW binding sites on PfHSP70-2 and PfHSP70-x determined from experimental data are shown in bold. Binding sites for other homologs predicted by sequence alignment are also listed.

| **PfHSP70 homolog** | **ADP binding residues** | **HEW binding residues** |
| --- | --- | --- |
| **PfHsp70-1** | Asp21, Gly23, Thr24, Tyr26, Thr25, Gly213, Gly214, Gly242, Glu281, Lys284, Arg285, Ser288, Gly351, Arg355, Ile356, Asp379 | Arg83, His100, Gly215, His239, Glu243, Asp244 |
| **PfHsp70-x** | Asp39, Gly41, Thr42, Thr43, Tyr44, Gly231, Gly232, Gly260, Glu299, Lys302, Arg303, Ser306, Gly369, Gly370, Ser371, Arg373, Ile374, Asp397 | **Arg101, His118, Gly233, His257, Glu261, Asp262** |
| **PfHsp70-2** | **Gly36, Thr37, Thr38, Tyr39, Gly223, Gly224, Gly252, Glu290, Lys293, Arg 294, Ser297, Gly360, Gly361, Arg364, Ile365, Asp388** | Arg96, Leu113, Gly225, His243, Glu253, Asp254 |
| **PfHsp70-3** | Gly10, Thr11, Thr12, Tyr13, Gly197, Gly198, Gly226, Glu264, Lys267, Arg268, Gly334, Gly335, Arg338, Ile339, Asp362, Ser271 | Arg70, Leu87, Gly199, His223, Glu227, Asp228 |

**Figure 2:**
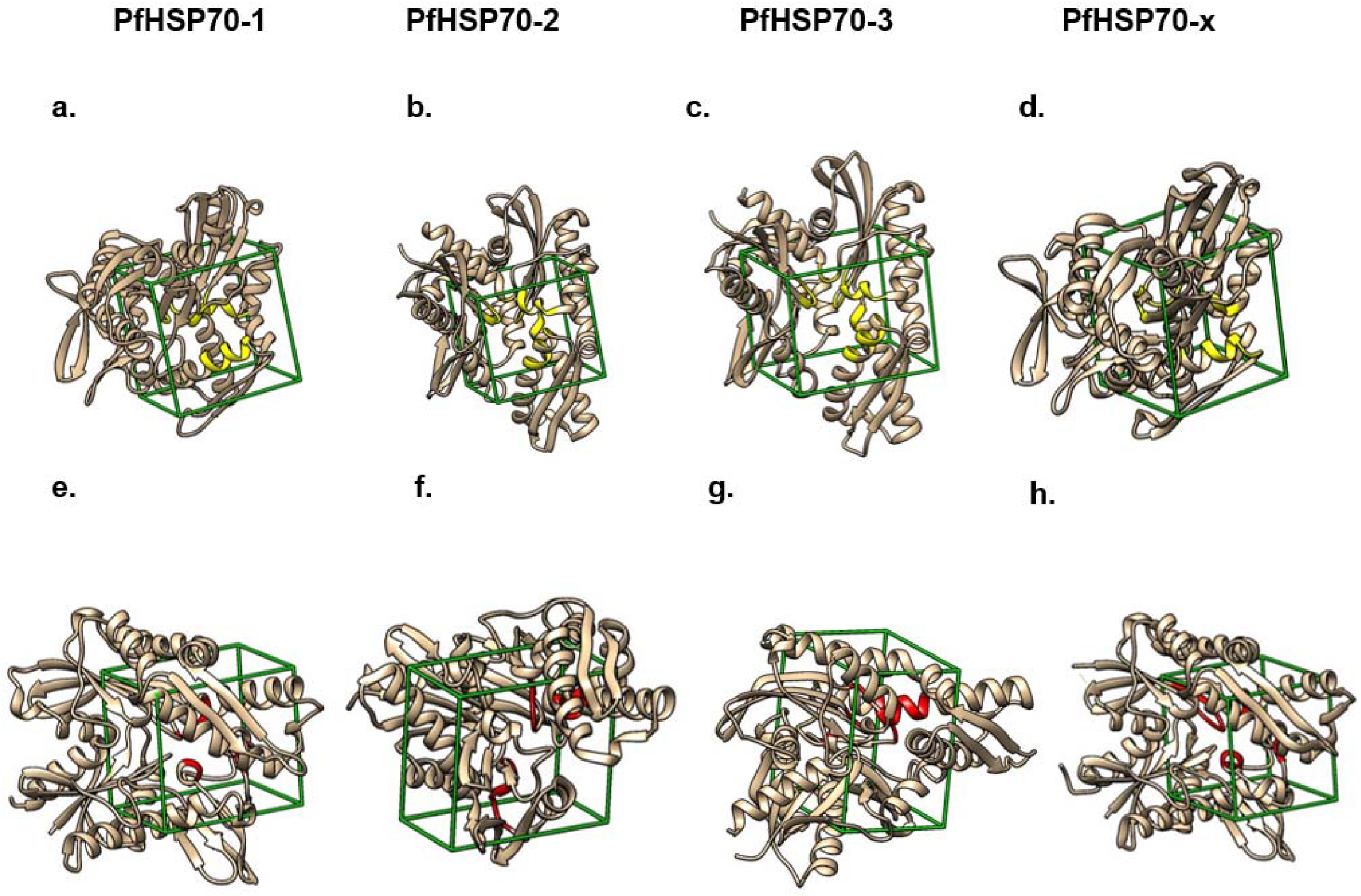
Grid boxes for molecular docking. Grid boxes (green) marked around ADP binding sites (yellow) (a-d) and HEW binding sites (red) (e-h). Protein molecules are shown as beige ribbon diagrams.

### Identification of ligand interacting residues

The binding energies obtained from the docking experiment for each pose were studied to select one best binding pose for each binding site on all PfHSP70s (Fig. 3). MKT-077 can be seen to be nestled into the respective binding pockets on each of the PfHSP70 proteins (Fig. 4). Binding energies for the selected poses are tabulated (Table 3). Analysis of these data has led us to understand that PfHSP70-2 and 3 are likely to bind with high affinity to MKT-077 (binding energies -7.38 and -7.356 respectively). PfHSP70-1 was predicted to bind the ligand with the least affinity and a significantly lower magnitude of binding energy (−6.118).

**Table 3:** The binding energies of MKT-077 with PfHSP70s at ADP and HEW binding sites. Binding energies were determined from Chimera after docking. Binding energy/ score were obtained from Chimera post molecular docking in Vina Plugin.

| **Protein name** | **ADP** | **HEW** |
| --- | --- | --- |
| **PfHsp70-1** | -6.118 | -5.94 |
| **PfHsp70-x** | -6.461 | -6.884 |
| **PfHsp70-2** | -7.38 | -6.398 |
| **PfHsp70-3** | -7.356 | -6.685 |

**Figure 3:**
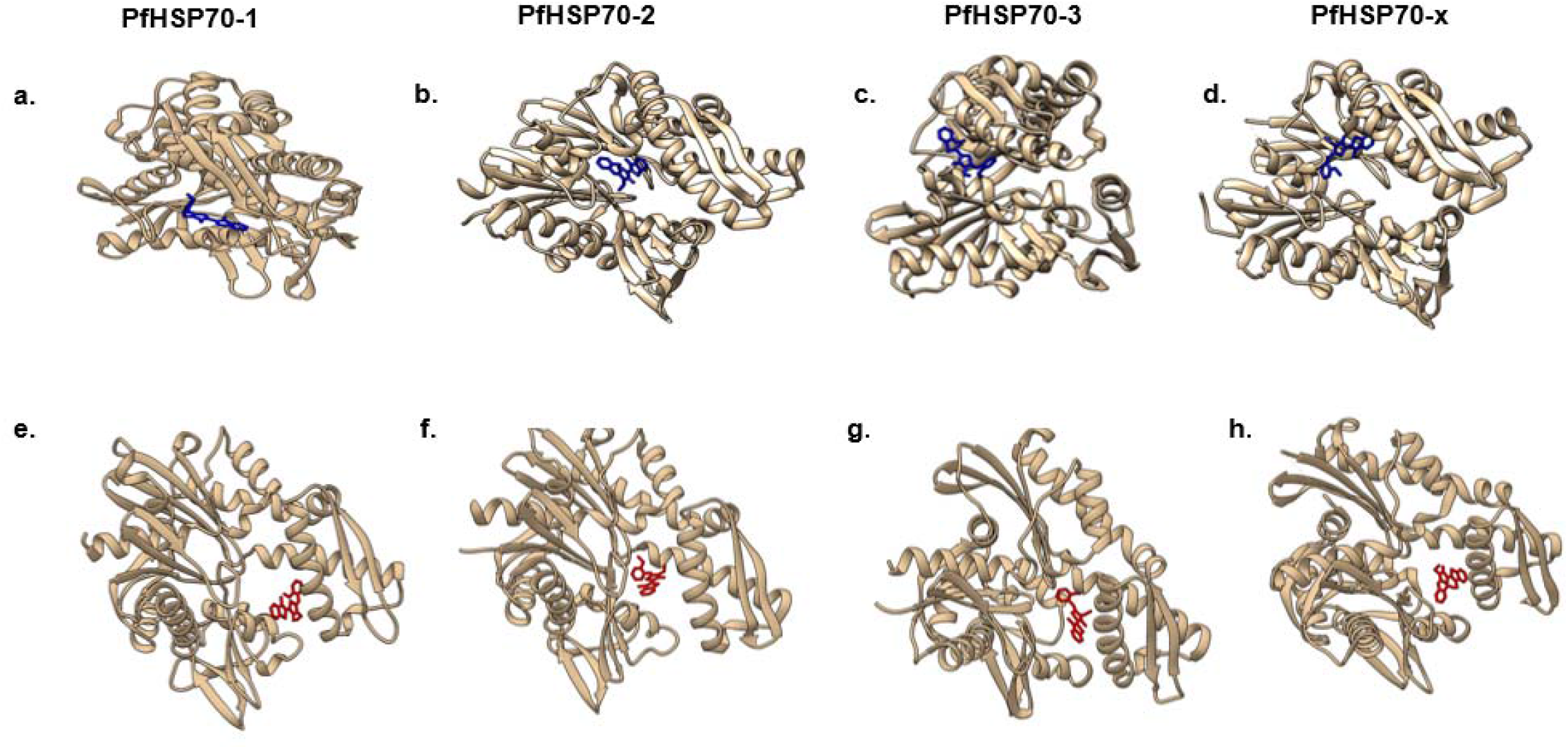
PfHSP70-MKT-077 complexes. MKT-077 docked on different PfHSP70 homolog (as labelled). Protein molecules are shown as beige ribbon diagrams a-d: MKT-077 (blue) docked at ADP binding site. e-h: MKT-077 (red) docked at HEW binding site.

**Figure 4:**
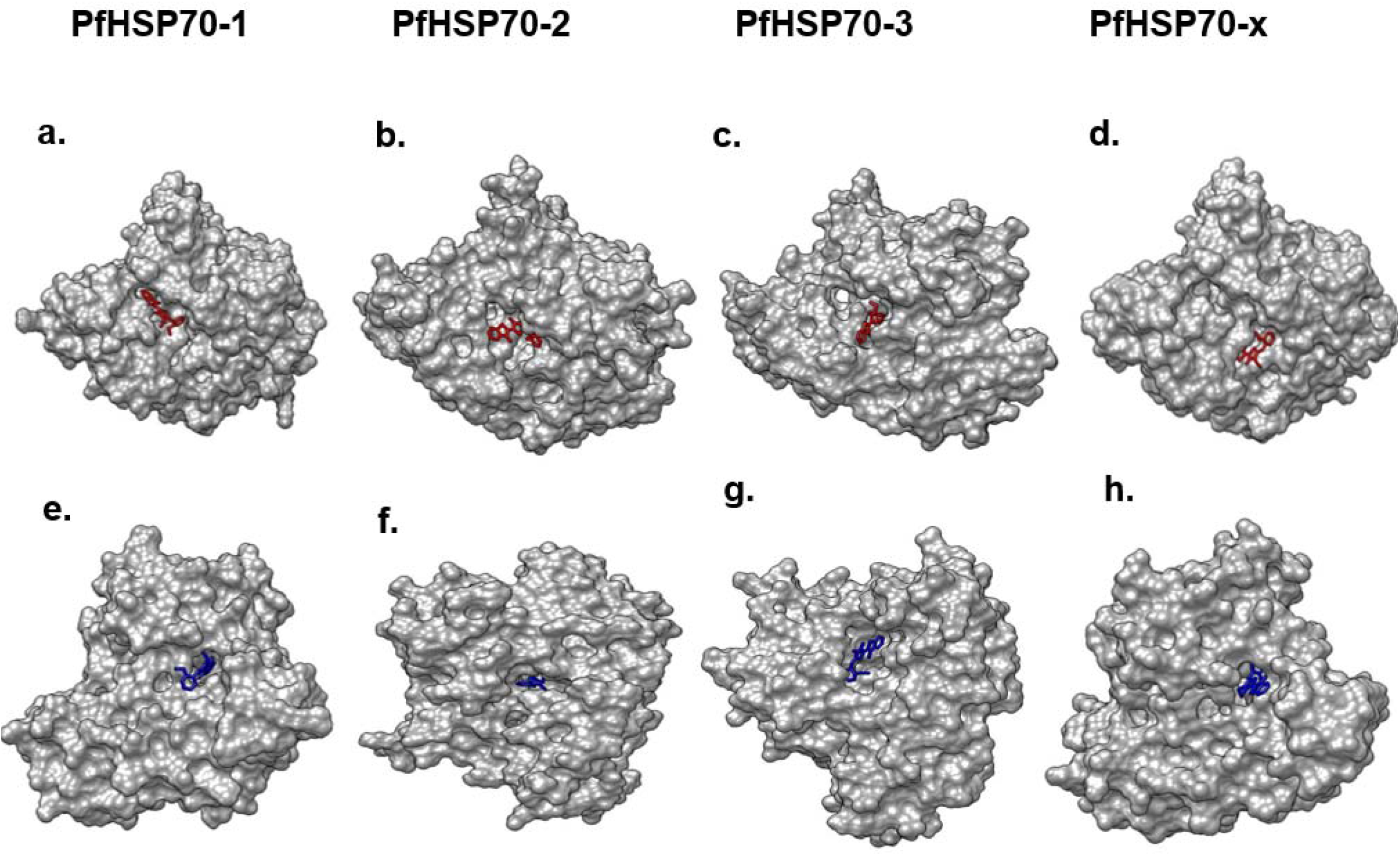
Molecular surface views of PfHSP70-MKT-077 complexes. a-d: MKT-077 (red) docked at ADP binding site of respective PfHSP70 homologs (as labelled). e-h: MKT-077 (blue) docked at HEW binding site of respective PfHSP70 homologs (as labelled).

We further analyzed the docked structures of PfHSP70-2 and 3 with MKT-077 to identify specific residues of these proteins that are involved in ligand binding by using LigPlot (Table 4). The interaction of MKT-077 with these PfHSP70 homologs is predicted to be mainly hydrophobic in nature (Fig. 5). Amino acids involved in these interactions mainly include Tyr, Ile, Arg and Asp (Table 4).

**Table 4:** MKT-077 binding residues on PfHSP70-2 and 3 as predicted by LigPlot analysis of docked complexes.

| **Protein name** | **MKT-077 interacting residues** |
| --- | --- |
| **PfHsp70-2** | Tyr 39, Asn59, Arg60, Ile61, Pro63, Glu77, Ile291, Arg294, Arg364, Asp388 |
| **PfHsp70-3** | Tyr 13, Ile 35, Gly 198, Arg 268, Ser 271, Gly 335, Arg 338, Asp 362 |

**Figure 5:**
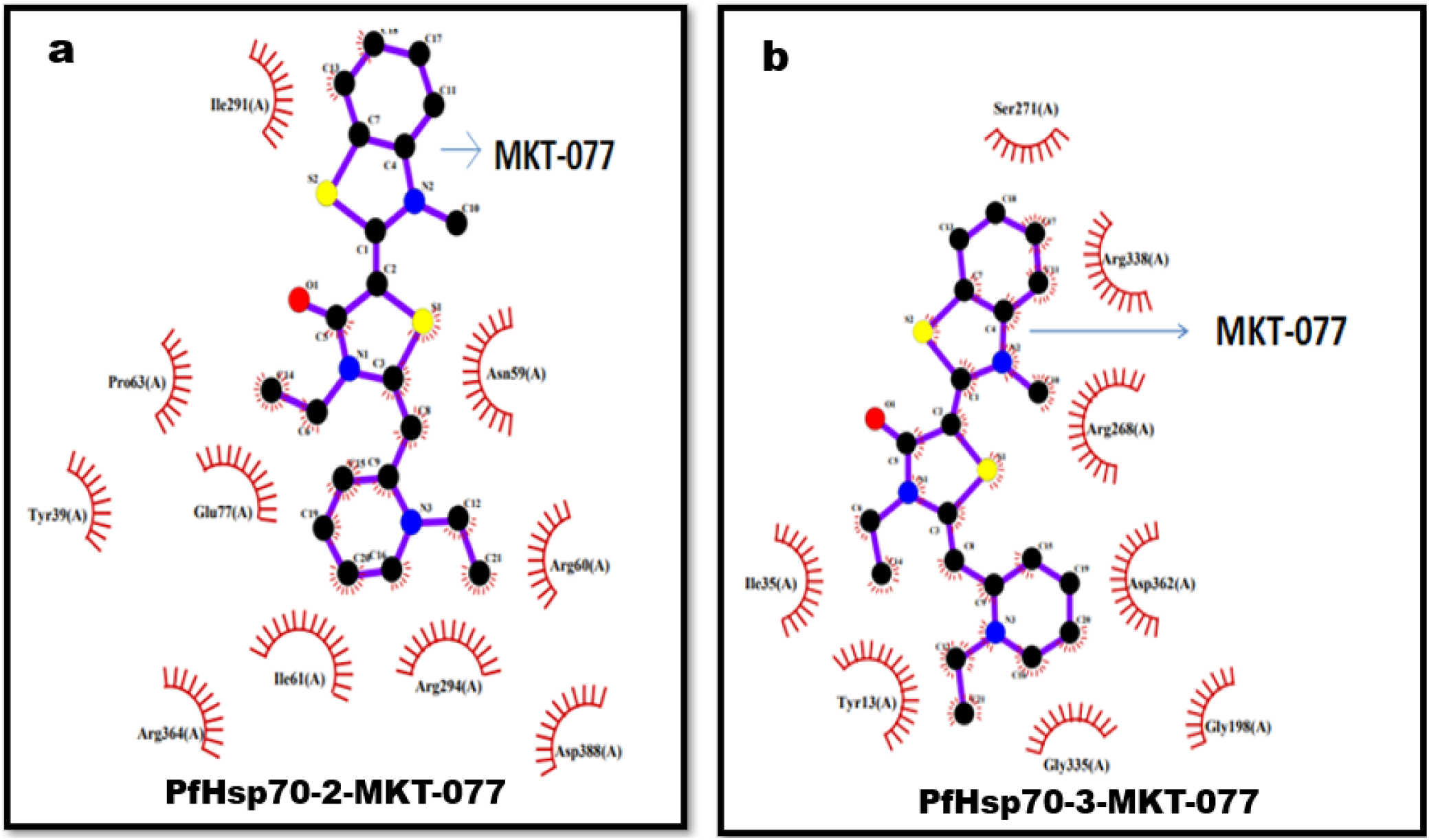
Ligplot analysis of complexes with best binding energy for. (a) PfHsp70-2-MKT-077 (b) PfHSP70-3-MKT-077 at ADP binding sites. MKT-077 is shown in the centre as ball and stick representation. Amino acids of PfHSP70s involved in hydrophobic interactions are depicted as arcs with emanating rays.

## Conclusion

Pf has four distinct HSP70 proteins that localize differently within parasites and infected erythrocytes. MKT-077 is a small molecule inhibitor that had shown immense potential in treatment of various diseases including malaria. It allosterically binds NBD of HSP70s and inhibits its ATPase activity. Reports on the ability of MKT-077 to inhibit *in vitro* growth of *Plasmodium* parasites led us to analyze binding of this inhibitor with PfHSP70s at the molecular level. Using *in silico* methods, we have predicted that MKT-077 is most likely to bind PfHSP70-1, 2 and 3 on its ADP binding site. On the other hand, the probability of binding of MKT-077 on PfHSP70-x is higher at the HEW binding site. The ER expressed PfHSP70-2 and the mitochondrial PfHSP70-3 are predicted to interact with MKT-077 with highest affinity. We therefore analyzed the structures of these two homologs complexed with MKT-077 to predict specific amino acid residues that are involved in the interactions. Information about specific interacting residues may form the basis for design of more effective and specific rhodacyanine drugs against malaria.

## Abbreviations

Pf: *Plasmodium falciparum*
HSP: heat shock protein
SBD: substrate binding domain
NBD: nucleotide binding domain
ATP: adenosine tri phosphate
ADP: adenosine di phosphate

## Acknowledgments

CN and VU were receiving scholarship from DBT, Government of India. SK is a DBT-SRF. The laboratories of PCM and RH are funded by UGC-DAE, and were earlier funded by DBT, RUSA and DST, Government of India.

## Declaration

*The authors have no relevant financial or non-financial interests to disclose*.

The authors declare that there are no conflicts of interest.

*All authors read and approved the final manuscript*.

## Supplementary Material

**Figure S1:**
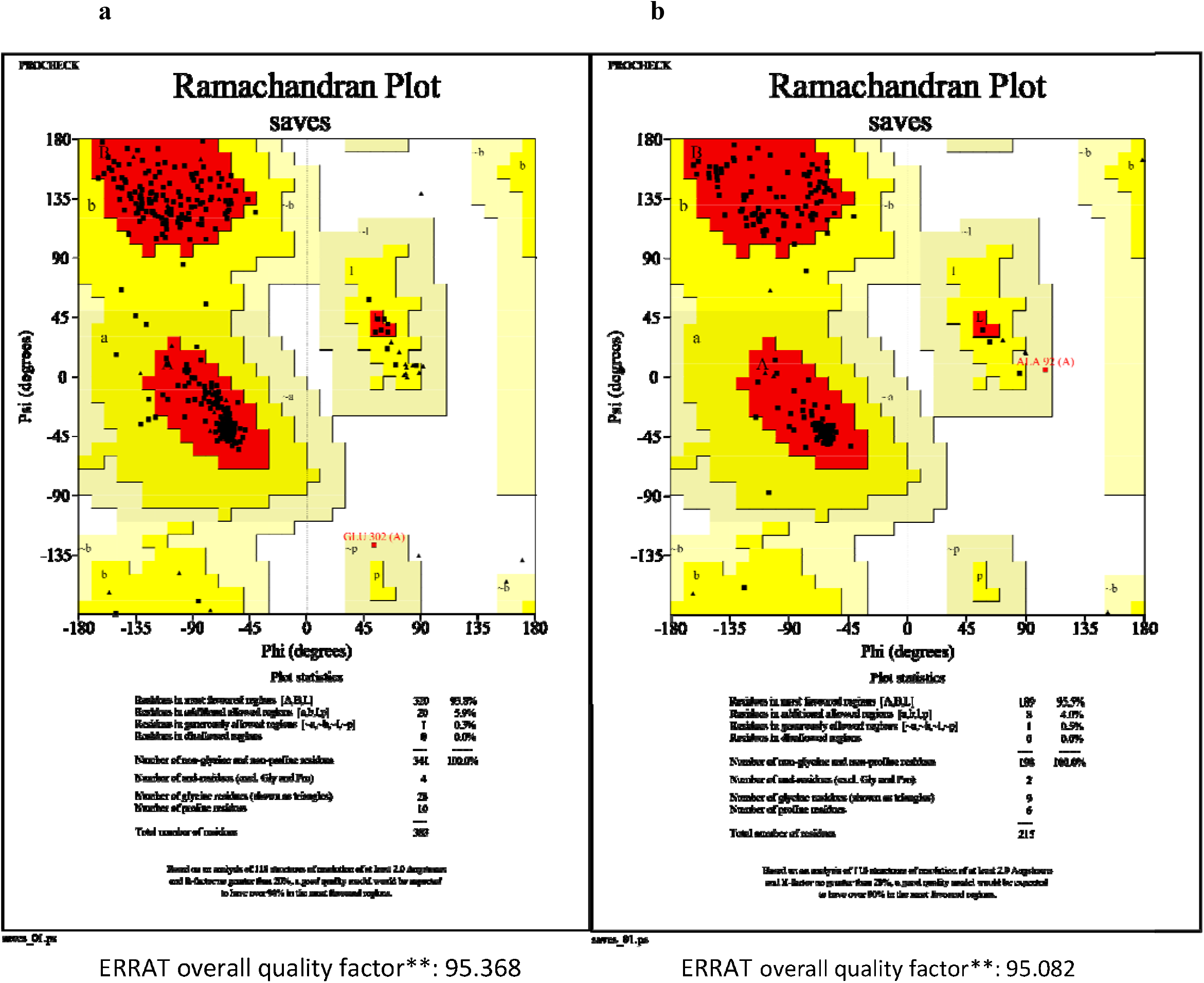
Validation of modeled structures. Ramachandran plot analysis of three dimensional structures obtained from SwissModel for a. PfHSP70-1 and b. PfHSP70-3. The overall quality factors obtained from ERRAT for each model are mentioned below the plot.

**Table S1:**
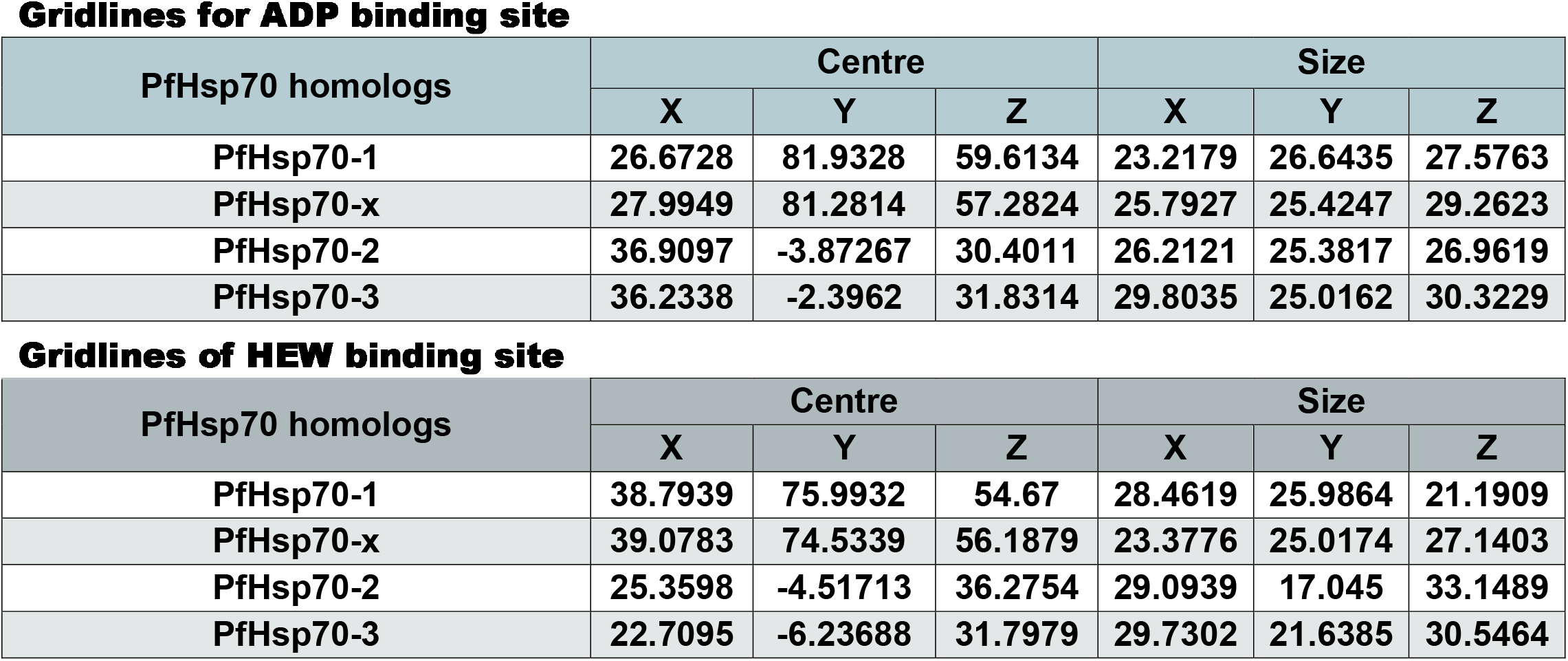
Grid box coordinates and dimensions. Tables shows center coordinates and size of the selected grid box for ADP and HEW binding sites on PfHSP70 homologs, as mentioned.

**Gridlines for ADP binding site**
| PfHsp70 homologs | Centre |  |  | Size |  |  |
| --- | --- | --- | --- | --- | --- | --- |
|  | X | Y | Z | X | Y | Z |
| PfHsp70-1 | 26.6728 | 81.9328 | 59.6134 | 23.2179 | 26.6435 | 27.5763 |
| PfHsp70-x | 27.9949 | 81.2814 | 57.2824 | 25.7927 | 25.4247 | 29.2623 |
| PfHsp70-2 | 36.9097 | -3.87267 | 30.4011 | 26.2121 | 25.3817 | 26.9619 |
| PfHsp70-3 | 36.2338 | -2.3962 | 31.8314 | 29.8035 | 25.0162 | 30.3229 |

| PfHsp70 homologs | Centre |  |  | Size |  |  |
| --- | --- | --- | --- | --- | --- | --- |
|  | X | Y | Z | X | Y | Z |
| PfHsp70-1 | 38.7939 | 75.9932 | 54.67 | 28.4619 | 25.9864 | 21.1909 |
| PfHsp70-x | 39.0783 | 74.5339 | 56.1879 | 23.3776 | 25.0174 | 27.1403 |
| PfHsp70-2 | 25.3598 | -4.51713 | 36.2754 | 29.0939 | 17.045 | 33.1489 |
| PfHsp70-3 | 22.7095 | -6.23688 | 31.7979 | 29.7302 | 21.6385 | 30.5464 |

